# Predicting Anorexia Nervosa and Autistic Characteristics in Individuals with Anorexia Nervosa from Resting State Cyclic Connectivity

**DOI:** 10.1101/2023.12.11.571092

**Authors:** Daniel Halls, Jenni Leppanen, Steve Williams, Kate Tchanturia

## Abstract

**Background:** Resting state functional Magnetic Resonance Imaging (rsfMRI) differences have been reported in individuals with anorexia nervosa (AN). However, methodological issues limit inferences, the dynamic cyclic nature of the resting state signal has not been explored, neither has the association with known elevated autistic characteristics.

**Methods:** 92 participants, 65 Individuals with AN and 27 controls underwent rsfMRI. Cyclic analysis was conducted to obtain pairwise relationships between regions. Classification of group and regression to predict autistic characteristics within the AN group was conducted, with model weights being explored to ascertain the most predictive pairwise relationships.

**Results:** Pairwise relationships in the temporal, dorsal and ventral attention networks were most predictive of group. The anterior intraparietal sulcus, salience, dorsal attention, auditory dorsal posterior cingulate cortex, and ventral attention networks were most predictive of autistic characteristics.

**Discussion:** Several distinct pairwise relationships predicting group and autistic characteristics were found, however, the global disruption of the temporal ordering of the cyclic wave, and variation in temporal ordering across resting state scan are also a neurophenotype in individuals with AN and the relationship to autistic characteristics. Characteristics associated with AN and autism are also predicted by distinct neural regions.

## 1. Introduction

Anorexia Nervosa (AN) is a life-threatening eating disorder (ED), characterised by intense concerns regarding weight gain, disturbance of body perception and restriction of dietary intake (American Psychiatric Association, 2013). The neurobiology of AN is poorly understood (Frank, Shott & DeGuzman, 2019), however the use neuroimaging, particularly resting state functional magnetic resonance imaging (rsfMRI) offers promise in demonstrating neurophenotypes that characterise AN. Previous work, utilising rsfMRI, has characterised atypical connectivity in individuals with AN involving temporal regions (Olivo et al., 2018), as well as the ventral attention, default mode and the salience networks (Spalatro et al., 2019; Gondo et al., 2023). Atypical connectivity, as demonstrated by rsfMRI, is also not just isolated to the underweight stage of illness course (Boehm et al., 2016; Gondo et al., 2023). Cross-sectional studies with weight restored individuals with AN has suggested altered connectivity involving the frontoparietal network (Boehm et al., 2016). Studies examining pre and post treatment in individuals with AN has also shown atypical connectivity involving the default mode and salience networks (Gondo et al., 2023).

A confounding difficulty when attempting to explore neurophenotypes in individuals with AN, is the presence autistic characteristics. Autism is defined as a neurodevelopmental disorder characterised by difficulties in social communication, restricted patterns of behaviour and repetitive interests (American Psychiatric Association, 2013). Over recent years the link between AN and autism has demonstrated on a neurobiological and behavioural level (Leslie et al., 2020; Tchanturia, Smith, Glennon, Burhouse, 2020; Kerr-Gaffney et al., 2021; Halls et al., 2022; Tchanturia, 2022). Behavioural work has shown that autistic characteristics are overrepresented in individuals with AN (Tchanturia et al., 2020; Kerr-Gaffney et al., 2021) and are associated with worse treatment outcomes (Leppanen, Sedgewick, Halls & Tchanturia, 2022). Previous work exploring the neurobiological link between autistic characteristics and AN is limited but, has either focused on structural similarities (Halls et al., 2022) or using task-based fMRI (Leslie et al., 2020). Despite the plethora of rsfMRI demonstrating atypical functional connectivity in autistic individuals (Reiter et al., 2021; Feng, M., & Xu, J 2023; Zhuang et al., 2023), no work to the best of our knowledge has explored autistic characteristics in individuals with AN using rsfMRI.

Previous work employing rsfMRI to explore neurophenotypes in AN demonstrate considerable drawbacks. This is because a majority of rsfMRI previous work done in individuals with AN uses “traditional” rsfMRI analysis (i.e., conducting correlation of time series from regions and running parametric/non-parametric models for significance testing) to explore neurophenotypes. A considerable drawback to this method is interpreting exactly what rsfMRI results mean. This is because in absence of an experimental stimuli it is difficult to know if group-based results represent meaningful neurophenotypes of illness or mental state (Gonzalez-Castillo, Kam, Hoy & Bandettini., 2021). Traditional rsfMRI that attempts to draw these types of inferences often uses logically invalid informal reverse inferencing, by affirming the consequent that brain region X will selectively represent a neurophenotypes of illness or mental state (Poldrack, 2006).

One potential solution to known reverse inferences challenges in rsfMRI analysis is the use of machine learning ([ML]; Zhuo, Li, Lin, Jiang, Xu, Tian, Wang & Song 2020). ML methods allow for valid formal reverse inferencing from neurobiology to group classification or mental state (Poldrack, 2011), as they are data driven approaches interested in the relationship between label and features (Khosla, Jamison, Ngo, Kuceyeski & Sabuncu, 2019). Arguably this makes ML methods more interpretable when exploring neurophenotypes (Abraham et al., 2017) than “traditional” rsfMRI methods. rsfMRI connectivity is also well suited to be explored using ML methods (Shahsavarani et al., 2020; Reiter et al., 2021), with it being used widely in rsfMRI analysis in autism (Reiter et al., 2021; Zhuang et al., 2023) and tinnitus research (Zimmerman, Abraham, Schmidt, Baryshnikov & Husain, 2018; Shahsavarani et al., 2020).

The second drawback of traditional rsfMRI used in previous work in AN, is traditional fMRI explores the synchronous nature of the neural resting state signal but ignores the cyclic dynamic nature of the signal. The resting state signal is known to be dynamic, as the spatio-temporal pattern of neural networks undergoes several reconfigurations during individual rsfMRI scan (Gonzalez-Castillo et al., 2021), due to traveling slow cyclic cortical waves (Baryshnikov & Schlafly 2016; Abraham, Shahsavarani, Zimmerman, Husain & Baryshnikov, 2021). This has been established experimentally from multichannel electroencephalography recordings that demonstrate propagating macroscopic slow cortical waves (Muller, Chavane, Reynolds & Sejnowski, 2018). This wave is cyclic but not periodic, meaning it is invariant to time (Baryshnikov & Schlafly 2016) and has an inherent temporal ordering (Abraham et al., 2021).

Interest in how the cyclicity and the temporal ordering differs in illness state when compared to healthy controls (HC) has over the past few years led for a search for methods to accurately explore these dynamic changes (Baryshnikov & Schlafly 2016; Abraham et al., 2021). Traditional correlation analysis has drawbacks as it assumes non-autocorrelated time series and is unable to capture temporal ordering (Shahsavarani et al., 2020). Granger causality assumes stationarity in the time series, which is rarely met, and dynamic time wrapping demonstrates poor reliability when compared to other functional connectivity measures (Shahsavarani et al., 2020). Finally other dynamic measures such as sliding window relies heavily on a priori filtering and hyperparameters (Baryshnikov & Schlafly, 2016). This led to the development of cyclic analysis (Baryshnikov & Schlafly, 2016). Cyclic analysis derives pairwise temporal relationships between time series using iterated path integrals, to find a lead-follower relationship between these pairwise time series as well as the temporal ordering of the cortical wave (Baryshnikov & Schlafly, 2016; Zimmerman et al., 2018; Shahsavarani et al., 2020; Abraham et al., 2021). Cyclic analysis has advantages over previous methods when exploring rsfMRI signal, for example it does not assume the rsfMRI signal to be stationary (Shahsavarani et al., 2020) and can recover the temporal ordering of the signal (Abraham et al., 2021). Cyclic analysis is also well suited for ML methods, as it produces a lead-lag matrix of temporal ordering which can easily be used in ML models (Zimmerman et al., 2018; Shahsavarani et al., 2020), and has been shown to generate more robust features than other measures such as dynamic time wrapping (Shahsavarani et al., 2020).

Therefore, this paper will use ML methods to classify the dynamic cyclic changes of the resting state signal obtained through cyclic analysis from individuals with AN and HC, to discover neurophenotypes in individuals with AN. The relationship between autistic characteristics and the temporal ordering of the underlying cyclic wave in individuals with AN will also be explored using ML methods, to address lack of such work in the field. Neurophenotypes will be explored by examining the model weights from the ML algorithms, to demonstrate the most predictive network, the most predictive pairwise relationship within that network as well as the most predictive between networks pairwise relationships.

## 2. Methods

### 2.1 Participants

A total of 93 participants, 66 individuals with AN and 27 HC were recruited from a wider longitudinal neuroimaging study (see Halls et al., 2022; Halls et al., 2023 and Leslie et al., 2020 for further details). Participants were all female, able to undergo an MRI scan with no history of brain injuries, neurological and learning disabilities. Before enrolment in the longitudinal study, participants underwent the Structured Clinical Interview for DSM-5 – research version (First et al., 2015) and gave written informed consent about study participation. Participants with AN were recruited from a variety of clinical services including the South London and Maudsley specialist ED Service, South-West London and St George’s ED Service as well as the UK ED charity Beat. All participants with AN had a diagnosis of AN, defined by the DSM-5 criteria, and were all at various illness stages, ranging from being acutely underweight to recovered. HC participants were recruited from the local community and King’s College London, and had no history of ED.

### 2.2 Behavioural Measures and analysis

Before the rsfMRI session participants completed the Hospital Anxiety and Depression scale (HADS; Zigmond & Snaith, 1983) as well as the Eating Disorders Examination-Questionnaire (EDE-Q; Fairburn & Beglin, 2008). These items exhibited acceptable internal consistency (Cronbach alpha = 0.73). Participants also provided information about their age, as well as height and weight to calculate body mass index (BMI, defined as kg/m^2^). To test for group differences, multiple bayesian regression with group as the X variable and the EDE-Q global score, age, BMI, anxiety, and depression as the y variables in each of the models. Weakly informative priors were obtained by scaling to the data (see supplementary material section 1 for further details). The means, standard deviations and high-density intervals are presented and were used as evidence for potential group difference. One HC was removed in the behavioural analysis due to being an outlier in EDE-Q global score (z score >3).

Assessing autistic characteristics was done in two ways. Participants with AN completed the Autistic Quotient 10 (Allison, Auyeung & Baron-Cohen, 2012), which was done just before the rsfMRI scan. Participants with AN also underwent the Autism Diagnostic Observation Schedule second edition (ADOS-2) to assess for autistic characteristics by a trained ADOS-researcher. This data was collected on average 2 years before the rsfMRI scan, as part of a longitudinal study (see Halls et al., 2021 and Leslie et al., 2020 for further details). The total score was used as a measure of autistic characteristics. In total 38 AN participants underwent the ADOS-2.

### 2.3 Scan acquisition

Images were acquired at the Centre for Neuroimaging Sciences at King’s College London on a 3T GE MRI scanner. A T1 weighted image was taken with the following parameters: echo time 3.02 seconds, repetition time 7.31 seconds, flip angle 11 degrees, field of view 270 mm, 256 x 256-pixel matrix and slice thickness of 1.2 mm. Three multi-echo images rsfMRI images were taken over 6.4 minutes period with 2 seconds repetition time, slice thickness 3 mm, slice thickness 3.3 mm, field of view 240 mm, 64 x 64-pixel matrix and a flip-angle 80 degrees.

### 2.5 MRI Preprocessing

Image quality was visually assessed using the automated quality control pipeline MRIQC (Esteban et al., 2017) and preprocessing was conducted using the standardised pre- processing pipeline fMRIPrep (Esteban et al., 2019, 2020; see supplementary material section 2 for further details). One AN participant’s scan was removed due to image acquisition artefacts, leaving the final sample at 92 individuals, (AN = 65, HC = 27) Following pre-processing, the images were detrended, band pass filtered at 0.01 - 0.08 Hetz, confounds removed and then standardised in Nilearn (https://nilearn.github.io/stable/index.html). The confounds chosen for removal was done using a data driven approach, by removing the number of aCompCors that explain 50% of the variance, along with the 24 movement regressors of: head motion, derivatives, squares, and squares of the derivatives of motion regressors (Mascali et al., 2021).

### 2.6 MRI analysis

#### 2.6.1 Seed Regions

39 regions of interest (ROI) were selected, taken from the multi-subject dictionary learning (MDSL) probabilistic atlas (Varoquaux, Gramfort, Pedregosa, Michel & Thirion, 2011) from Nilearn. The MDSL atlas has been shown to superior to other atlas for classification tasks (Abraham et al., 2017). The time series extracted from these ROIs by taking the average across all the voxels in the ROIS then being standardized to zero mean and scaled to unit variance. The ROIs, the co-ordinates, and the functional network that the region belongs to are listed in table 1.

**Table 1:** Table of region of interests, networks, and co-ordinates.

| Labels | Network | Co-ordinates (X, Y, Z) |
| --- | --- | --- |
| Left auditory | Auditory | -53.28, -8.88, 32.36 |
| Right auditory | Auditory | 53.47, -6.49, 27.52 |
| Striate | Striate | 1.18, -74.54, 10.79 |
| Left default mode network | Default mode network | -45.8, -64.78, 31.84 |
| Medial default mode network | Default mode network | -0.2, -55.21, 29.87 |
| Front default mode network | Default mode network | -0.15, 51.42, 7.58 |
| Right default mode network | Default mode network | 51.66, -59.34, 28.88 |
| Occipital posterior | Occipital posterior | 0.41, -91.05, 1.58 |
| Motor | Motor | -1.48, -27.93, 61.5 |
| Right dorsolateral prefrontal cortex | Right ventral attention | 40.1, 20.96, 44.72 |
| Right front pole | Right ventral attention | 37.83, 55.49, 1.22 |
| Right parietal | Right ventral attention | 47.53, -52.42, 43.06 |
| Right posterior temporal | Right ventral attention | 62.53, -32.99, -9.14 |
| Basal | Basal | -0.91, -2.75, 6.15 |
| Left parietal | Left ventral attention | -41.66, -59.04, 44.61 |
| Left dorsolateral prefrontal cortex | Left ventral attention | -39.04, 19.28, 43.27 |
| Left frontal pole | Left ventral attention | -40.08, 50.65, 0.81 |
| Left intraparietal sulcus | Dorsal attention | -29.39, -59.43, 44.2 |
| Right intraparietal sulcus | Dorsal attention | 31.6, -58.09, 45.69 |
| Left lateral occipital complex | Visual sec | -30.54, -85.14, 9.1 |
| Visual | Visual sec | -24.29, -74.28, -11.74 |
| Right lateral occipital complex | Visual sec | 33.4, -77.96, 4.31 |
| Dorsal anterior cingulate cortex | Salience | -28.17, 46.32, 21.56 |
| Ventral anterior cingulate cortex | Salience | -0.45, 34.06, 20.73 |
| Right anterior insula | Salience | 28.38, 47.72, 22.13 |
| Left superior temporal sulcus | Temporal | -52.12, -17.92, 13.28 |
| Right superior temporal sulcus | Temporal | 52.61, -13.65, 12.11 |
| Left temporoparietal junction | Language | -55.52, -43.77, 10.08 |
| Broca | Language | -48.66, 25.11, 5.7 |
| Superior frontal sulcus | Language | -3.39, 17.19, 63.52 |
| Right temporoparietal junction | Language | 54.42, -29.5, -2.72 |
| Right pars opercularis | Language | 52.38, 29.39, 2.93 |
| Cerebellum | Cerebellum | 1.05, -58.49, -23.91 |
| Dorsal posterior cingulate cortex | Dorsal posterior cingulate cortex | -1.44, -59.12, 55.25 |
| Left insula | Cingulate-Insula | -41.33, 13.63, 2.7 |
| Cingulate | Cingulate-Insula | 1.05, 9.2, 46.43 |
| Right insula | Cingulate-Insula | 43.01, 14.3, 2.79 |
| Left anterior intraparietal sulcus | Anterior intraparietal sulcus | -47.85, -32.07, 41.9 |
| Right anterior intraparietal sulcus | Anterior intraparietal sulcus | 48.36, -29.04, 43.13 |

#### 2.6.2 Cyclic Analysis

It is beyond the scope of this paper to give a full account of cyclic analysis, for further details on the method, please see Abraham et al., 2021. Cyclic analysis was performed on each participant’s time series to generate a 39x39 lead-lag matrix, using python. Each value in the lead lag matrix represents the strength of the relationship between time series and the sign indicating direction of information flow from nodes, with zero representing no cyclic relationship (Shahsavarani et al., 2020).

Once the lead-lag matrix was built, the cyclic ordering of the propagating wave was recovered from participants with AN and HC. This can be done as the order of the components of the eigenvector correspond to the leading eigenvalue of the lead matrix, indicating cyclic order of the propagating wave (Abraham et al., 2021). Therefore, the eigenvectors and eigenvectors from the lead-lag matrices of all participants was calculated, as well as calculating a mean temporal ordering of AN and HC participants, by extracting the eigenvectors and eigenvectors from a mean lead-lag matrix for each group.

For the ML models, the lower triangle of the 39x39 lead-lag matrices of each participant were extracted and the diagonals removed, as to remove repeated and uninformative information. The 2D matrices were reshaped into a single dimensional vector, before being concatenated into a single 2D matrix with each row representing a participant and each column a pairwise relationship between regions, which was used as the X variable in all the models.

#### 2.6.3 Classification models

Several classification model algorithms were built to discover neurophenotypes underlying AN. Models were scored by accuracy and area under the receiver operating curve (AUC ROC) using stratified cross-validation, which randomly samples from the minority and majority classes to the original distribution, so each fold has the same distribution as the initial distribution, achieving more robust validation (Szeghalmy & Fazekas, 2023). As our classes for group classification where imbalanced (HC = 27, AN = 65), which is known to cause bias (Teh, Armitage, Tesfaye, Selvarajah & Wilkinson 2020), the oversampling method assembled synthetic minority oversampling technique was used. This technique has been shown to be suitable for rsfMRI and superior to no oversampling technique in predicting accuracy (Teh et al., 2020). As the primary aim of this study is to explore which features are most relevant in classifying AN and HC scans and not necessarily building the most accurate model, the main criteria for model selection was to have interpretable explorable weights, with a regularization parameter. The models compared were a random forest, support vector machine classifier, a logistic regression and a stacked model of a random forest and logistic regression. For all models, the X variable was the concatenated lead lag cyclic vectors of all participants, while the y was a categorical label indicating which group the vector belonged to. The model algorithm that performed the best was chosen as the final model (see supplementary material section 3 for further details). The stacked model performed the best (*accuracy = 0.87, AUC ROC = 0.96*) and was used as the final model.

To explore neurophenotypes underlying AN, the most predictive network with the most predictive pairwise relationship between regions within that network, as well as the most predictive pairwise relationship between networks regions from the model, were extracted. This was done by converting the logistic regression coefficients and random forest feature importance into z scores, to allow meaningful comparison between the logistic regression coefficients and feature importance. The maximum z score for mean network values, within and between regions was obtained to decide which temporal ordering was most predictive. The maximum z score was converted to a bayes factor (BF) to quantify the strength of evidence that regions and the network are the most predictive.

Finally, the pairwise time series from the most predictive within and between network regions was explored. This involved calculating the pairwise accumulated oriented area from the time series of the most predictive pairwise relationships (see Abraham et al., 2021 for further details). The relationship was calculated for each participant, then average was taken for HC and participants with AN.

#### 2.6.4 Predicting autistic characteristics

To explore the relationship between autistic characteristics and rsfMRI within the AN group, regression models predicting the ADOS total score and AQ10 were built, with sample sizes of 38 and 65 respectively. A random forest, as well as lasso, ridge and support vector regression models were used, to ascertain which model was best able to predict autistic characteristics, as these models have a regularization parameter and interpretable weights. Scoring for all models was done by examining r2 and mean absolute error (MAE). For the AQ10 models scoring was done by stratified cross validation, however scoring for the ADOS models, due to the small sample size causing class imbalance, was done using a train and hold out dataset consisting of 30 and 8 participants respectively. Initially for all models the X matrix was every pairwise relationship within the lead lag matrix from each participant, however, this resulted in all the models performing arbitrarily badly (see supplementary material section 4). Filtering features from the lead lag matrix, by retaining the number of connections that exhibited a small Pearson r correlation (0.1) to the AQ10 and ADOS total scores, did improve model fit greatly to acceptable levels (see supplementary material section 5 for further details). The support vector regression model was the best fit for the AQ10 (*r2 = 0.33, MAE = -1.58*) and for consistency a support vector regression was also used as the final model for the ADOS total score (*r2 = 0.41, MAE = 6.58*). Like the classification strategy, the most predictive network, regions within the most predictive network and the most predictive between network regions was examined by finding the maximum support vector regression coefficient.

#### 2.6.4 Cyclicity and behavioural measures

To explore the relationship between cyclicity and behavioural measures, correlation of behavioural measures to the cyclic ordering as well as pairwise relationships of most predictive lead lag relationships was conducted. The EDE-Q global score, depressive and anxiety traits (from the HADS), age and BMI were correlated to the variation in participants leading eigenvector component phase, which determines cyclic ordering. This was done using Pearson’s r in the HC and AN groups, with confidence intervals (CI), r value and BF used to assess strength of evidence. The variation in the pairwise accumulated oriented area of most predictive pairwise relationship was also correlated by group to the EDE-Q global score, depressive and anxiety traits (from the HADS), age and BMI using the same strategy.

## 3. Results

### 3.1 Clinical measures

Individuals with AN had increased EDE-Q, anxiety, and depression scores with lower BMI than HC controls. No group differences were suggested for age (see table 2).

**Table 2.** Summary of Bayesian Regression Models and Behavioural Measures.

| Measure | Mean | SD | HDI 3% | HDI 97% |
| --- | --- | --- | --- | --- |
| EDE-Q scores [HC] | -1.68 | 0.32 | -2.26 | -1.06 |
| Anxiety scores [HC] | -5.78 | 0.93 | -7.49 | -4.01 |
| Depression scores [HC] | -3.97 | 0.96 | -5.75 | -2.18 |
| BMI [HC] | 5.26 | 0.90 | 3.65 | 7.00 |
| Age [HC] | 0.40 | 0.80 | -0.11 | 1.92 |
|  | AN mean (SD) |  | HC mean (SD) |  |
| EDE-Q scores | 2.55 (1.54) |  | 0.87 (0.60) |  |
| Anxiety scores | 11.78 (4.22) |  | 6.00 (3.25) |  |
| Depression scores | 6.89 (4.32) |  | 2.92 (2.90) |  |
| BMI | 19.48 (3.30) |  | 24.75 (5.00) |  |
| Age | 21.68 (3.54) |  | 22.07 (3.08) |  |
| Abbreviations: AN: Anorexia Nervosa; BMI: Body Mass Index; EDE-Q: Eating Disorders Examination-Questionnaire; HDI: High Density Interval; SD: Standard Deviation |  |  |  |  |

### 3.3 Cyclicity order

Table 3 and figure 1 demonstrates the cyclic ordering of the propagated wave in the AN and HC groups.

**Figure 1:**
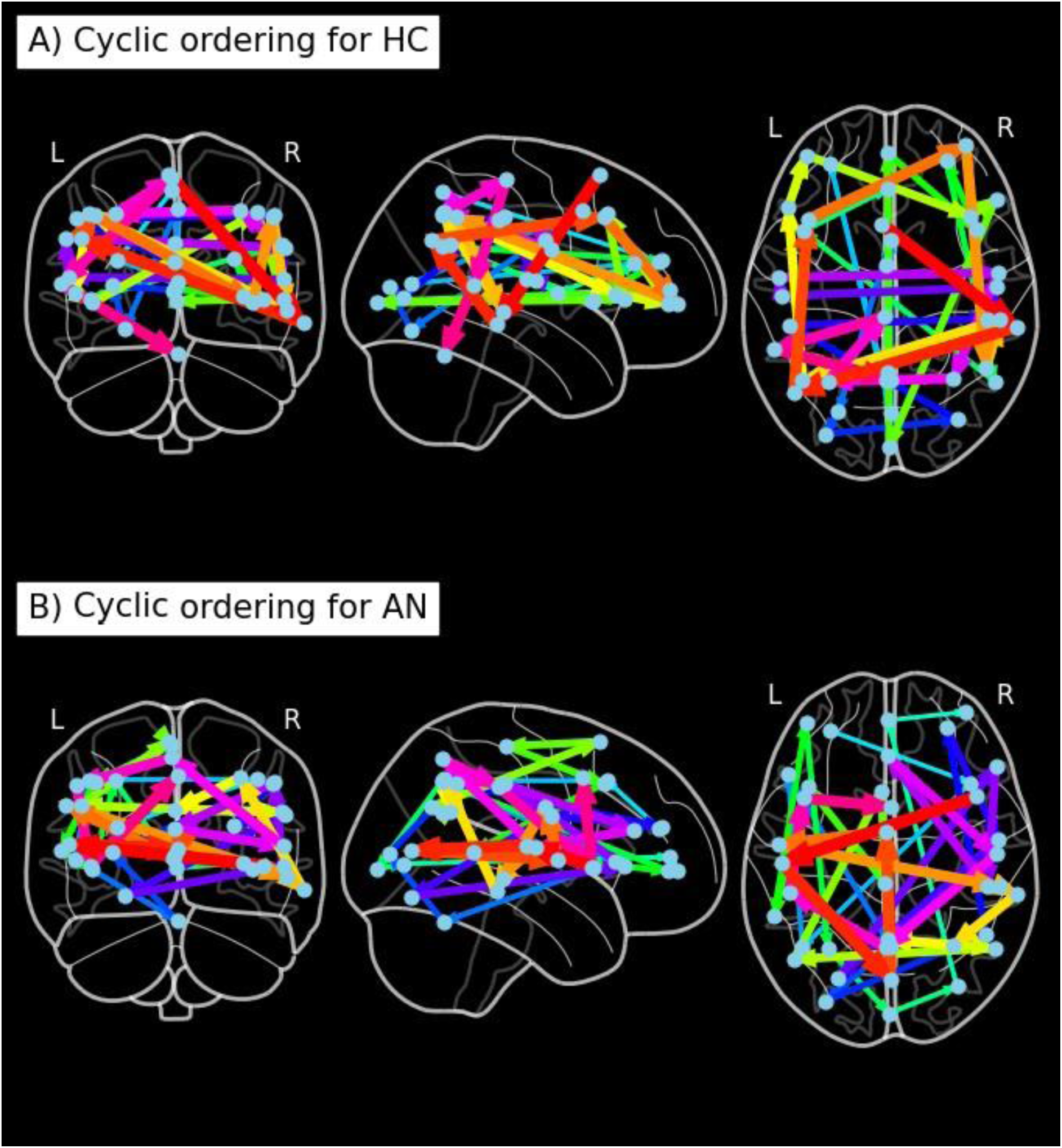
Temporal Ordering of the Cyclic Wave for A) Healthy Controls, B) Individuals with Anorexia Nervosa

**Table 3.** Cyclic ordering by Group.

| Anorexia Nervosa participants Cyclic Ordering | Healthy controls Cyclic Ordering |
| --- | --- |
| Right insula<br>Left superior temporal sulcus<br>Striate<br>Basal<br>Left auditory<br>Right Temporo-parietal junction<br>Right posterior temporal<br>Right intraparietal sulcus<br>Medial default mode network<br>Right default mode network<br>Left default mode network<br>Superior front sulcus<br>Motor<br>Left dorsolateral prefrontal cortex<br>Left temporo-parietal junction<br>Left front pole<br>Left intraparietal sulcus<br>Occipital posterior<br>Right lateral occipital cortex<br>Front default mode network<br>Right front pole<br>Dorsal anterior cingulate cortex<br>Right dorsolateral prefrontal cortex<br>Left parietal<br>Broca<br>Cerebellum | Superior frontal sulcus<br>Right posterior temporal<br>Left default mode network<br>Left dorsolateral prefrontal cortex<br>Right front pole<br>Right parietal<br>Right temporo-parietal junction<br>Left parietal<br>Broca<br>Left front pole<br>Right dorsolateral prefrontal cortex<br>Right pars opercularis<br>Occipital posterior<br>Front default mode network<br>Right insula<br>Right anterior insula<br>Right default mode network<br>Left insula<br>Ventral anterior cingulate cortex<br>Basal<br>Medial default mode network<br>Dorsal posterior cingulate cortex<br>Dorsal anterior cingulate cortex<br>Striate<br>Cingulate<br>Visual |
| Left lateral occipital cortex<br>Right parietal<br>Right anterior Insula<br>Right auditory<br>Visual<br>Right pars opercularis<br>Right anterior intraparietal sulcus<br>Ventral anterior cingulate cortex<br>Right superior temporal sulcus<br>Dorsal posterior cingulate cortex<br>Left anterior intraparietal sulcus<br>Left Insula<br>Cingulate | Left lateral occipital cortex<br>Right lateral occipital cortex<br>Left Ant intraparietal sulcus<br>Right Ant intraparietal sulcus<br>Right superior temporal sulcus<br>Left superior temporal sulcus<br>Left auditory<br>Right auditory<br>Right intraparietal sulcus<br>Left intraparietal sulcus<br>Motor<br>Left temporo-parietal junction<br>Cerebellum |

### 3.4 Classification of rsfMRI

A stacked model of random forest and logistic regression was able to classify with 86% accuracy, AN and HC rsfMRI cyclic connectivity with an AUC ROC = 0.97. The most predictive network for classifying scans was the temporal network, with strong evidence that this was the most predictive network (see table 4 and figure 2). Within the temporal network the pairwise relationship between the right and left superior temporal sulcus (see table 4 and figure 2) was the most predictive (due to this being the only within network pairwise relationship). For individuals with AN, this relationship was weaker than in HC, and was from left superior temporal sulcus to right, while in HC the relationship was reversed. The most predictive between network relationship for classifying AN and HC scans was the left intra-parietal sulcus to the right post temporal gyrus, representing a relationship from the right ventral attention network to the dorsal attention network (see table 4 and figure 2). This relationship was weaker for individuals with AN than HC, with the relationship going from the left intra-parietal sulcus (dorsal attention network) to the right post temporal gyrus (right ventral attention network) while a vice versa relationship was observed for HCs.

**Figure 2:**
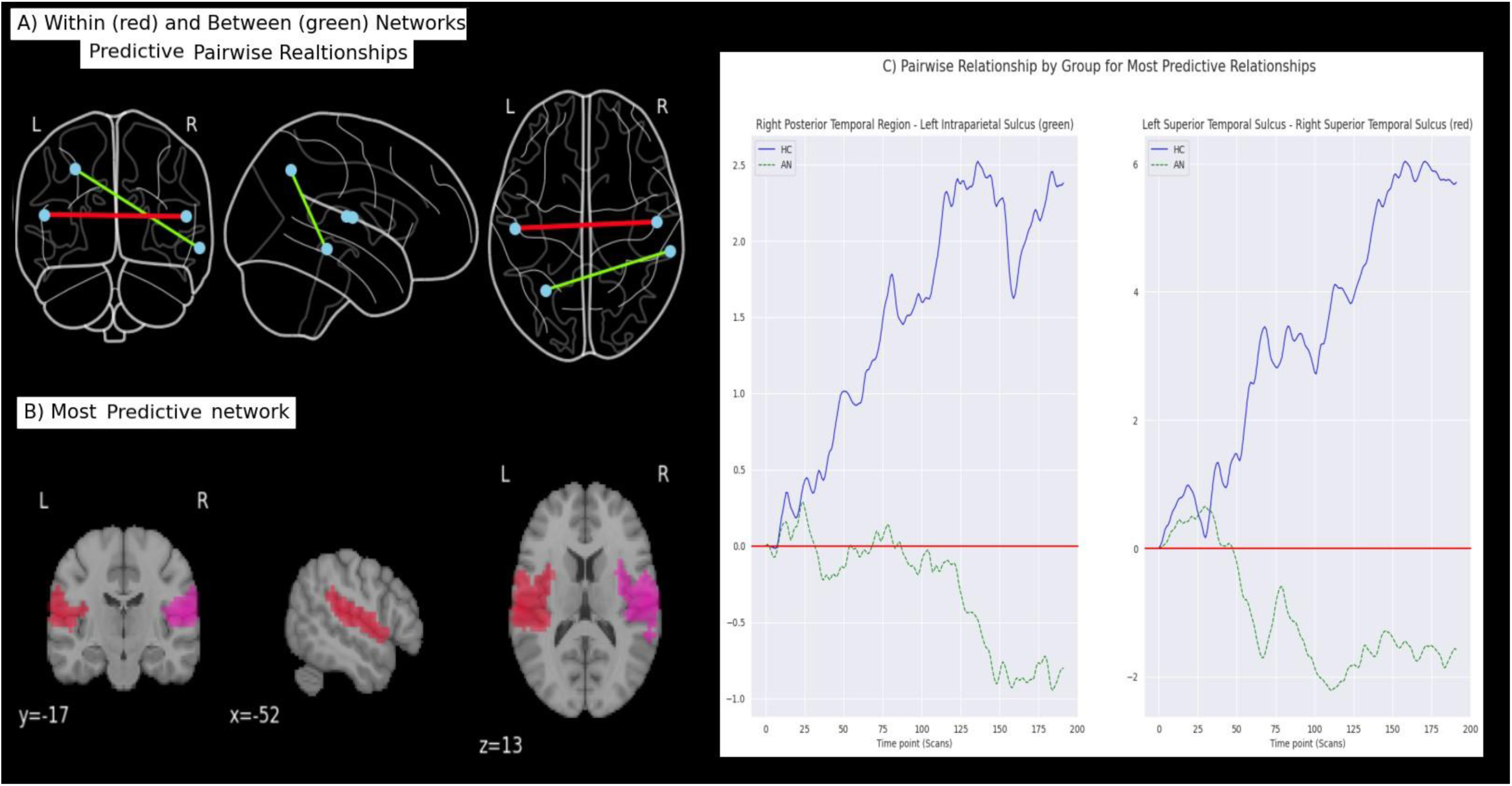
Most Predictive Pairwise Relationships for Group Classifications: A) Within (red) and Between (green) Networks; B) Most Predictive Network, C) Temporal ordering of most Predictive Relationships

**Table 4.** Predictive regions and networks from the classification model, classifying AN and HC from rsfMRI.

|  |  |  |  |  |  |  |
| --- | --- | --- | --- | --- | --- | --- |
| Most Predictive Network |  |  |  |  |  |  |
| Network | RF importance | Log Coefficients | Maximum Z score | Lead lag AN | Lead lag HC | BF |
| Temporal | 0.01 | -0.01 | 2.83 | 0.79 | -2.37 | 54.87 |
| Most Predictive within network regions |  |  |  |  |  |  |
| Regions | RF importance | Log Coefficients | Maximum Z score | Lead lag AN | Lead lag HC | BF |
| R STS - L STS | 0.01 | -0.01 | N/A* | 0.79 | -2.37 | N/A* |
| Most predictive between networks regions |  |  |  |  |  |  |
| Regions (network) | RF importance | Log Coefficients | Maximum Z score | Lead lag AN | Lead lag HC | BF |
| L IPS - R Post Temp<br>(D Att - R V Att) | 0.03 | -0.02 | 11.45 | 1.60 | -5.71 | >100 |
| *The temporal network only has one lead lag relationship in it, therefore calculating z scores and BF is not possible. |  |  |  |  |  |  |
| Abbreviations: AN: Anorexia Nervosa; BF: Bayes Factor; D Att: Dorsal Attention Network; HC: Healthy Controls; L: Left; Log: Logistic Regression; Post Temp: Posterior Temporal; R: Right; RF: Random Forest; STS: Superior Temporal Sulcus; V Att: Ventral Attention Network. |  |  |  |  |  |  |

### 3.5 Predicting of Autistic characteristics from rsfMRI

The most predictive network for autistic characteristics as measured by the AQ10 was the anterior intraparietal sulcus network, with the most predictive within region relationship for AQ10 models being the temporal ordering of right to the left anterior intraparietal sulcus. Increasing autistic characteristics suggested a strengthening of this relationship. The most predictive between network relationship for autistic characteristics as measured by the AQ10 was from the left intra-parietal sulcus to the right auditory region, representing a pairwise relationship from the dorsal attention to the auditory network. Increasing autistic characteristics suggested this relationship was strengthened (see table 5 and figure 3).

**Figure 3:**
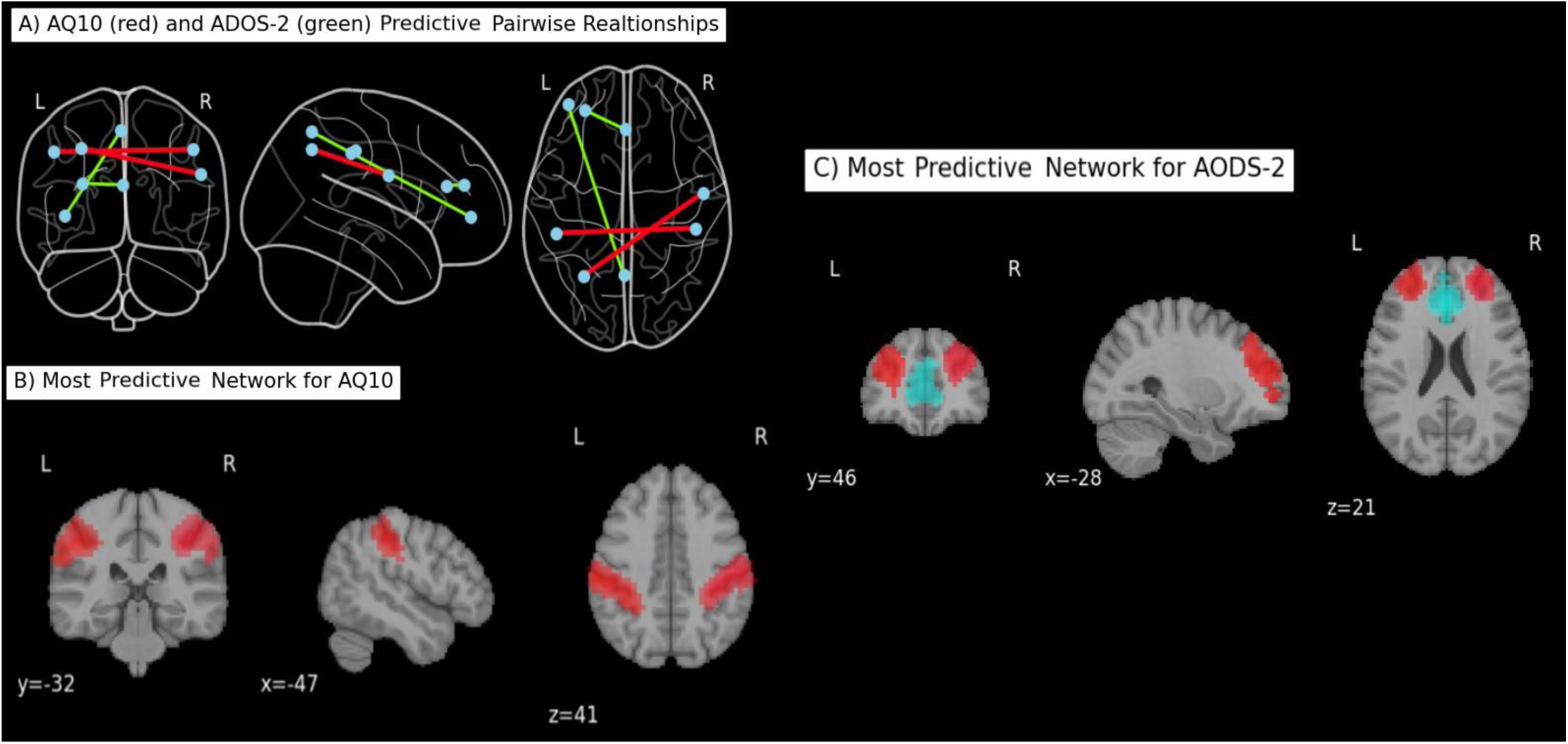
Most Predictive Pairwise Relationships for Predicting Autistic Characteristics: A) AQ10 (red) and ADOS-2 (green); B) Most Predictive Network for AQ10; C) Most Predictive Network for ADOS-2

**Table 5.**
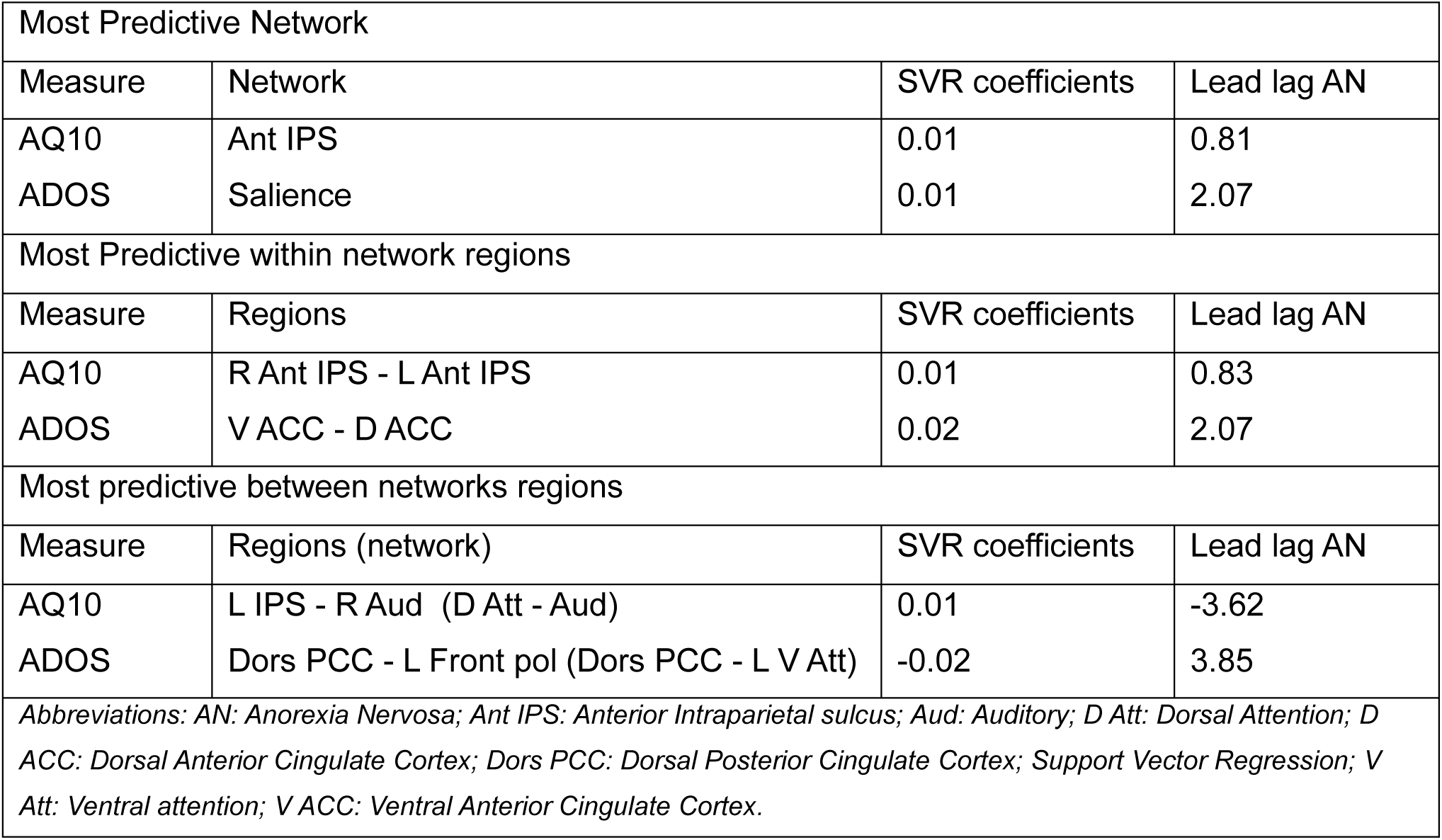
Predictive regions and networks from the regression model predicting autistic characteristics from rsfMRI.

For the ADOS-2 model the most predictive network was the salience network, with the most predictive within network pairwise relationship being from the dorsal to ventral anterior cingulate cortex. Increasing autistic characteristics suggested an increasing relationship from the left to the right anterior intraparietal sulcus. The most predictive between network pairwise relationship was from the dorsal posterior cingulate cortex (dorsal posterior cingulate network) to the left frontal pole (left ventral attention network). Increasing autistic characteristics suggested a weakening of this relationship (see table 5 and figure 3).

### 3.6 Cyclicity and behavioural measures

A moderate correlation, but with weak evidence, in the HC group of the variation in the pairwise accumulated oriented area of the left intra-parietal sulcus to the right post temporal gyrus was found to EDE-Q global score (*r = 0.44, BF = 2.66, 95% CI [0.06, 0.71]*). No other correlations with sufficient evidence existed in either group (see supplementary material section 6).

Both depression (*r = 0.30, BF = 2.63, 95% CI [0.06, 0.51]*).) and anxiety (*r = 0.26, BF = 1.30, 95% CI [0.02, 0.48]*) traits were weakly correlated to the cyclic ordering in the AN group with weak evidence. No other correlations with sufficient evidence existed in either group (see supplementary material section 7).

## 4. Discussion

By using ML and novel cyclic analysis, we have explored the cyclic dynamic resting state changes seen in AN and the relationship with autistic characteristics in this group. We have demonstrated several important pairwise temporal relationships in AN and autistic characteristics seen in AN. Our work has also shown a further relationship between anxiety, depression, and ED symptoms and the resting dynamic cyclic wave.

Our first set of main findings were the neurophenotypes that underline of AN. Pairwise relationships between the right and left superior temporal sulcus in temporal network as well as the left intra-parietal sulcus (dorsal attention network) and right posterior temporal (right ventral attention network) were the most predictive neurophenotypes in AN. Putting these findings in context of previous work is difficult, as fundamentally this study is exploring different aspects of the resting state signal compared to previous work. However, our work does fit in with previous studies examining dynamic variation in the resting state signal in individuals with AN (Spalatro et al., 2019; Seidel et al., 2020). Increased neuronal variability in the ventral attention network in individuals with AN and individuals with bulimia nervosa when compared to HC has been reported (Spalatro et al., 2019). This neuronal variability was correlated to BMI, and ED characteristics (Spalatro et al., 2019). Differences in low frequency fluctuations involving regions in the temporal network has been observed in individuals with AN compared to HC (Seidel et al., 2020). Finally, group differences in the temporal network have been found when examining the synchronous nature of the neural resting state signal (Olivo et al., 2018). Taken with results from our study, it seems that resting dynamic and synchronous changes in the neural signal in the ventral attention and temporal networks are potential neurophenotypes in AN.

Our next set of findings was regarding the neurophenotypes of autistic characteristics in individuals with AN. Most predictive pairwise relationships where: the right and left anterior intraparietal sulcus (representing the anterior intraparietal sulcus network), the ventral and dorsal anterior cingulate (salience network), right auditory region and left anterior intraparietal sulcus (right auditory and dorsal attention network) and the dorsal posterior cingulate and left frontal pole (dorsal posterior cingulate cortex/left ventral attention network). Autistic characteristics increased all these temporal ordering apart from the dorsal posterior cingulate and left frontal pole ordering where it decreased this temporal ordering. Again, putting these findings in context of pervious work is difficult, as in AN to the best of our knowledge there are no rsfMRI studies exploring the link between autistic characteristics and AN, also cyclic analysis has not been conducted in autism. However, previous work using ML exploring dynamic functional connectivity in autistic individuals found that weights involving the dorsal and ventral attention networks were the most predictive of autistic characteristics measured by ADOS scores (Zhuang et al., 2023). Previous ML working exploring the synchronous changes in the neural signal has also indicated that the dorsal attention network as an important neurophenotype in autistic individuals (Reiter et al., 2021). Similar to this study, using the ADOS as a measure of autistic characteristics, the salience network has also been found to be significantly different between autistic individuals and non-autistic individuals (Yang et al., 2021). Taken with our work this suggests that dynamic and synchronous changes across a range of networks, including the dorsal/ventral attention networks as well as the salience network are important neurophenotypes for autistic characteristics, which is potentially shared in individuals with AN and autistic individuals.

Our results also suggest several interesting findings beyond predictive pairwise relationships, that are important regarding the neurophenotypes underlying individuals with AN. A potential mechanism for differences between groups in the pairwise relationship is the variability and ordering in that relationship over time. Over time the pairwise relationship between most predictive nodes varied between groups, in HCs the relationship became stronger with direction of flow static, while in individuals with AN, the direction of flow of the cyclic wave was more variable between nodes. Also interestingly, in the HC group ED traits were positively correlated with variability in the pairwise relationship. This suggests that ED traits in the HC group are associated with an increasingly variable pairwise relationship, becoming more “AN like”. Taken together, this perhaps suggests that the variability in the pairwise relationship is the salient feature in predicting group, however further work is needed to confirm this hypothesis.

There was little overlap in the most predictive pairwise relationships classifying group and predicting autistic characteristics. Though pairwise relationships in the dorsal attention network were involved in both classifying group and predicting autistic characteristics, the other pairwise relationships did not overlap, suggesting on a neural level a delineation of autistic and ED traits. Behavioural work exploring autistic characteristics in individuals with AN has found similar profiles in autistic characteristics, but with subtle differences, with some items of the ADOS adequately discriminate between individuals with AN and autistic individuals (Kerr-Gaffney et al., 2021). Our results may suggest how this behavioural finding is encoded on a neural level. Moreover, the AQ10 and the ADOS were predicted from different pairwise relationships. This could be due to methodological difficulties, especially with the ADOS-2, as the ADOS-2 models could not be cross-validated, had a smaller sample than the AQ10 models and was conducted two years earlier than the rsfMRI. However, it maybe that the ADOS-2 and AQ10 measure fundamentally different things. There is some suggestion that self-reported AQ10 may not entirely be measuring autistic characteristics. Previous work has found that the AQ10 is associated more strongly to obsessive-compulsive and anxiety and depressive traits in individuals with AN than the ADOS-2 (Leppanen et al., 2022). The AQ10 has also been criticised for its inability to distinguish between anxiety and autism (Ashwood et al., 2016). Our previous longitudinal work also highlighted the difficulties of using the AQ10 in a research setting (Halls et al., 2023). Though previous meta-analysis has found increased AQ10 scores in individuals with AN, the authors acknowledged that this differences could be due to co-morbidities such as anxiety (Westwood et al., 2016). It seems then our neurobiological findings may support previous behavioural work suggesting that the AQ10 and the ADOS-2 are measuring different psychological difficulties.

Finally, despite our findings demonstrating more predictive pairwise relationships for group and autistic characteristics, the model weights for these pairwise relationships are relatively small, with many pairwise relationships contribute to the models. Therefore, while some pairwise relationships are more important than others in classification and regression, global disruption in the cortical wave is likely also a neurophenotype in individuals with AN and autistic characteristics among individuals with AN. Previous work does support the hypothesis that global changes are an important neurobiological feature in individuals AN and autistic characteristics in this group. Global network alteration of the resting state signal has been reported in both autistic individuals (Xie et al., 2022) and individuals with AN (Geisler et al., 2019). In individuals with AN it was interesting that anxiety and depressive traits were associated with more variability in the global temporal ordering of this wave. Recent behavioural work has highlighted that the presence of anxiety and depression is associated with more severe ED symptoms (Sander, Moessner & Bauer, 2021). Therefore, a possible mechanism for this behavioural finding is that people with higher anxiety and depression traits, the temporal ordering of the dynamic cortical wave is more variable, which, our ML model have shown is predictive of having AN.

This study is not without its limitations. This study was unable to delineate AN participants by illness state due to sample size issues, resulting in the AN group being a heterogeneous mix of participants at differing illness state. This study took part in the COVID-19 pandemic which greatly impacted upon recruitment, meaning that we were unable to meaningfully separate AN participants based on illness stage due to sub-groups having too small a sample size. Also, regarding the AN group, this study did not have information on what type of treatment modality participants with AN were receiving or had received. All AN participants had treatment at some point in their illness journey, but this study did not have access to full medical records and participants were recruiting from a wide range of clinical services and a charity organisation (Beat). Finally, lack of ADOS-2 before the rsfMRI is a limitation, as though the ADOS-2 results have been shown to stable over time in measuring autistic characteristics (Bieleninik et al., 2017) an up-to-date ADOS-2 with a larger sample size would allow for more robust predictions from our autistic trait models.

In conclusion by using novel cyclic analysis and ML methods this study has demonstrated several neurophenotypes in individuals with AN and the relationship to autistic characteristics in this group. Our work suggests that variation in the temporal ordering of pairwise relationship is an important mechanism underlying the neurobiology of AN. But also, that the global disruption of temporal ordering of the dynamic wave is also a neurophenotype in individuals with AN. Our work also suggests that autistic characteristics and ED traits are distinct on a neural level. Importantly this work contributes to the emerging neurobiological literature exploring the link between ED and autism. Finally, our work has been able to demonstrate that novel cyclic analysis is an interesting method to explore the underlying neurobiology of individuals with AN and the relationship to autistic characteristics.

