## Supplementary Material for "Predicting Anorexia Nervosa and Autistic Characteristics in Individuals with Anorexia Nervosa from Resting State Cyclic Connectivity"

**Section 1: Priors for the Logistic Regression Models.**

EDE-Q global score ~ group

Family name: Gaussian

Link: identity

Observations: 89

Priors:

Common-level effects

Intercept ~ Normal(mu: 2.0764, sigma: 4.5243)

group ~ Normal(mu: 0, sigma: 8.5364)

Auxiliary parameters

sigma ~ HalfStudentT(nu: 4, sigma: 1.5346)

anxiety ~ group

Family name: Gaussian

Link: identity

Observations: 89

Priors:

Common-level effects

Intercept ~ Normal(mu: 10.1573, sigma: 13.8906)

group ~ Normal(mu: 0, sigma: 26.2087)

Auxiliary parameters

sigma ~ HalfStudentT(nu: 4, sigma: 4.7117)

depression ~ group

Family name: Gaussian

Link: identity

Observations: 89

Priors:

Common-level effects

Intercept ~ Normal(mu: 5.7753, sigma: 12.7303)

group ~ Normal(mu: 0, sigma: 24.0195

Auxiliary parameters

sigma ~ HalfStudentT(nu: 4, sigma: 4.3181)

BMI ~ group

Family name: Gaussian

Link: identity

Observations: 88

Priors:

Common-level effects

Intercept ~ Normal(mu: 21.0377, sigma: 13.4651)

group ~ Normal(mu: 0, sigma: 24.7723)

Auxiliary parameters

sigma ~ HalfStudentT(nu: 4, sigma: 4.5209)

age ~ group

Family name: Gaussian

Link: identity

Observations: 91

Priors:

Common-level effects

Intercept ~ Normal(mu: 21.7953, sigma: 9.9985)

group ~ Normal(mu: 0, sigma: 18.7055)

Auxiliary parameters

sigma ~ HalfStudentT(nu: 4, sigma: 3.3801)

**Section 2: fMRIPrep pre-processing**

Results included in this manuscript come from preprocessing performed using fMRIPrep 22.0.2 (Esteban, Markiewicz, et al. (2018); Esteban, Blair, et al. (2018); RRID:SCR_016216), which is based on Nipype 1.8.5 (K. Gorgolewski et al. (2011); K. J. Gorgolewski et al. (2018); RRID:SCR_002502).

Anatomical data preprocessing

A total of 1 T1-weighted (T1w) images were found within the input BIDS dataset.The T1-weighted (T1w) image was corrected for intensity non-uniformity (INU) with N4BiasFieldCorrection (Tustison et al. 2010), distributed with ANTs 2.3.3 (Avants et al. 2008, RRID:SCR_004757), and used as T1w-reference throughout the workflow. The T1w-reference was then skull-stripped with a Nipype implementation of the antsBrainExtraction.sh workflow (from ANTs), using OASIS30ANTs as target template. Brain tissue segmentation of cerebrospinal fluid (CSF), white-matter (WM) and gray-matter (GM) was performed on the brain-extracted T1w using fast (FSL 6.0.5.1:57b01774, RRID:SCR_002823, Zhang, Brady, and Smith 2001). Brain surfaces were reconstructed using recon-all (FreeSurfer 7.2.0, RRID:SCR_001847, Dale, Fischl, and Sereno 1999), and the brain mask estimated previously was refined with a custom variation of the method to reconcile ANTs-derived and FreeSurfer-derived segmentations of the cortical gray-matter of Mindboggle (RRID:SCR_002438, Klein et al. 2017). Volume-based spatial normalization to one standard space (MNI152NLin2009cAsym) was performed through nonlinear registration with antsRegistration (ANTs 2.3.3), using brain-extracted versions of both T1w reference and the T1w template. The following template was selected for spatial normalization: ICBM 152 Nonlinear Asymmetrical template version 2009c [Fonov et al. (2009), RRID:SCR_008796; TemplateFlow ID: MNI152NLin2009cAsym].

Functional data preprocessing

For each of the 1 BOLD runs found per subject (across all tasks and sessions), the following preprocessing was performed. First, a reference volume and its skull-stripped version were generated from the shortest echo of the BOLD run using a custom methodology of fMRIPrep. Head-motion parameters with respect to the BOLD reference (transformation matrices, and six corresponding rotation and translation parameters) are estimated before any spatiotemporal filtering using mcflirt (FSL 6.0.5.1:57b01774, Jenkinson et al. 2002). BOLD runs were slice-time corrected to 0.969s (0.5 of slice acquisition range 0s-1.94s) using 3dTshift from AFNI (Cox and Hyde 1997, RRID:SCR_005927). The BOLD time-series (including slice-timing correction when applied) were resampled onto their original, native space by applying the transforms to correct for head-motion. These resampled BOLD time-series will be referred to as preprocessed BOLD in original space, or just preprocessed BOLD. A T2 star map was estimated from the preprocessed EPI echoes, by voxel-wise fitting the maximal number of echoes with reliable signal in that voxel to a monoexponential signal decay model with nonlinear regression. The T2 star /S0 estimates from a log-linear regression fit were used for initial values. The calculated T2 star map was then used to optimally combine preprocessed BOLD across echoes following the method described in (Posse et al. 1999). The optimally combined time series was carried forward as the preprocessed BOLD. The BOLD reference was then co-registered to the T1w reference using bbregister (FreeSurfer) which implements boundary-based registration (Greve and Fischl 2009). Co-registration was configured with six degrees of freedom. First, a reference volume and its skull-stripped version were generated using a custom methodology of fMRIPrep. Several confounding time-series were calculated based on the preprocessed BOLD: framewise displacement (FD), DVARS and three region-wise global signals. FD was computed using two formulations following Power (absolute sum of relative motions, Power et al. (2014)) and Jenkinson (relative root mean square displacement between affines, Jenkinson et al. (2002)). FD and DVARS are calculated for each functional run, both using their implementations in Nipype (following the definitions by Power et al. 2014). The three global signals are extracted within the CSF, the WM, and the whole-brain masks. Additionally, a set of physiological regressors were extracted to allow for component-based noise correction (CompCor, Behzadi et al. 2007). Principal components are estimated after high-pass filtering the preprocessed BOLD time-series (using a discrete cosine filter with 128s cut-off) for the two CompCor variants: temporal (tCompCor) and anatomical (aCompCor). tCompCor components are then calculated from the top 2% variable voxels within the brain mask. For aCompCor, three probabilistic masks (CSF, WM and combined CSF+WM) are generated in anatomical space. The implementation differs from that of Behzadi et al. in that instead of eroding the masks by 2 pixels on BOLD space, a mask of pixels that likely contain a volume fraction of GM is subtracted from the aCompCor masks. This mask is obtained by dilating a GM mask extracted from the FreeSurfer’s aseg segmentation, and it ensures components are not extracted from voxels containing a minimal fraction of GM. Finally, these masks are resampled into BOLD space and binarized by thresholding at 0.99 (as in the original implementation). Components are also calculated separately within the WM and CSF masks. For each CompCor decomposition, the k components with the largest singular values are retained, such that the retained components’ time series are sufficient to explain 50 percent of variance across the nuisance mask (CSF, WM, combined, or temporal). The remaining components are dropped from consideration. The head-motion estimates calculated in the correction step were also placed within the corresponding confounds file. The confound time series derived from head motion estimates and global signals were expanded with the inclusion of temporal derivatives and quadratic terms for each (Satterthwaite et al. 2013). Frames that exceeded a threshold of 0.5 mm FD or 1.5 standardized DVARS were annotated as motion outliers. Additional nuisance timeseries are calculated by means of principal components analysis of the signal found within a thin band (crown) of voxels around the edge of the brain, as proposed by (Patriat, Reynolds, and Birn 2017). The BOLD time-series were resampled into standard space, generating a preprocessed BOLD run in MNI152NLin2009cAsym space. First, a reference volume and its skull-stripped version were generated using a custom methodology of fMRIPrep. All resamplings can be performed with a single interpolation step by composing all the pertinent transformations (i.e. head-motion transform matrices, susceptibility distortion correction when available, and co-registrations to anatomical and output spaces). Gridded (volumetric) resamplings were performed using antsApplyTransforms (ANTs), configured with Lanczos interpolation to minimize the smoothing effects of other kernels (Lanczos 1964). Non-gridded (surface) resamplings were performed using mri_vol2surf (FreeSurfer).

Many internal operations of fMRIPrep use Nilearn 0.9.1 (Abraham et al. 2014, RRID:SCR_001362), mostly within the functional processing workflow. For more details of the pipeline, see the section corresponding to workflows in fMRIPrep’s documentation.

Copyright Waiver

The above boilerplate text was automatically generated by fMRIPrep with the express intention that users should copy and paste this text into their manuscripts unchanged. It is released under the CC0 license.

**Section 3: Model Scoring Parameters for the Classification Models**

| Model | Accuracy | Area under the curve |
| --- | --- | --- |
| Support Vector Classifier | 0.72 | 0.90 |
| Decision tree | 0.73 | 0.73 |
| Random forest | 0.87 | 0.94 |
| Logistic | 0.83 | 0.94 |
| Stacked model | 0.87 | 0.96 |

**Section 4: Model Scoring Parameters for the Regression Models with no Feature Selection**

|  | Autism quotient 10 scores | | Autism Diagnostic Observation Schedule second edition | |
| --- | --- | --- | --- | --- |
| Model | R2 | Mean Absolute error | R2 | Mean Absolute error |
| Lasso regression | -0.035 | -2.073 | -0.464 | 9.770 |
| Ridge Regression | -0.303 | -2.337 | -0.061 | 8.639 |
| Random Forest | -0.003 | -2.088 | -0.292 | 9.397 |
| Support Vector Regression | -0.281 | -2.314 | -0.060 | 8.640 |

**Section 5: Model Scoring Parameters for the Regression Models with Feature Selection**

|  | Autism quotient 10 scores | | Autism Diagnostic Observation Schedule second edition | |
| --- | --- | --- | --- | --- |
| Model | R2 | Mean Absolute error | R2 | Mean Absolute error |
| Lasso regression | -0.09 | -2.081 | -0.464 | 9.770 |
| Ridge Regression | 0.321 | -1.583 | 0.410 | 6.557 |
| Random Forest | 0.084 | -2.008 | -0.225 | 9.095 |
| Support Vector Regression | 0.326 | -1.579 | 0.405 | 6.582 |

**Section 6: Correlation of Behavioural Measures to variation in the pairwise accumulated oriented areas of most predictive regions for group classification.**

| Measures | HC: R Post Temp - L IPS | HC: L STS - R STS | AN: R Post Temp - L IPS | AN: L STS - R STS |
| --- | --- | --- | --- | --- |
| EDE-Q Global | r: 0.44  CI95%: [0.06, 0.71]  BF10: 2.66 | r: -0.21  CI95%: [-0.55, 0.19]  BF10: 0.40 | r: -0.02  CI95%: [-0.26, 0.24]  BF10: 0.16 | r: -0.05  CI95%: [-0.29, 0.21]  BF10: 0.17 |
| Anxiety | r: -0.08  CI95%: [-0.45, 0.32]  BF10: 0.26 | r: -0.01  CI95%: [-0.39, 0.38]  BF10: 0.24 | r: -0.07  CI95%: [-0.31, 0.18]  BF10: 0.18 | r: 0.19  CI95%: [-0.06, 0.42]  BF10: 0.47 |
| Depression | r: 0.07  CI95%: [-0.33, 0.44]  BF10: 0.26 | r: -0.22  CI95%: [-0.56, 0.19]  BF10: 0.42 | r: 0.04  CI95%: [-0.22, 0.28]  BF10: 0.17 | r: 0.13  CI95%: [-0.12, 0.37]  BF10: 0.27 |
| Age | r: -0.15  CI95%: [-0.5, 0.24]  BF10: 0.31 | r: 0.03  CI95%: [-0.35, 0.4 ]  BF10: 0.24 | r: 0.01  CI95%: [-0.24, 0.26]  BF10: 0.16 | r: 0.01  CI95%: [-0.24, 0.26]  BF10: 0.16 |
| BMI | r: -0.09  CI95%: [-0.45, 0.30]  BF10: 0.26 | r: -0.09  CI95%: [-0.46, 0.3]  BF10: | r: 0.11  CI95%: [-0.15, 0.35]  BF10: 0.23 | r: -0.11  CI95%: [-0.35, 0.15]  BF10: 0.22 |
| *Abbreviations: AN: Anorexia Nervosa; BF10: Bayes Factor; BMI: Body Mass Index; CI95%: Confidence Interval; EDE-Q: Eating Disorders Examination- Questionnaire; HC: Healthy Controls; L: Left; Post Temp: Posterior Temporal; R: Right; STS: Superior Temporal Sulcus.* | | | | |

**Section 7: Correlation of Variation of the Eigenvector Component Phase to Behavioural Measures**

| Measures | HC eigenvector component phase | AN eigenvector component phase |
| --- | --- | --- |
| EDE-Q Global | r: -0.07  CI95%: [-0.44, 0.33]  BF10: 0.26 | r: 0.16  CI95%: [-0.10, 0.39]  BF10: 0.32 |
| Anxiety | r: -0.08  CI95%: [-0.45, 0.32]  BF10: 0.26 | r: 0.26  CI95%: [0.02, 0.48]  BF10: 1.30 |
| Depression | r: -0.19  CI95%: [-0.54, -0.21]  BF10: 0.37 | r: 0.30  CI95%: [0.06, 0.51]  BF10: 2.63 |
| Age | r: 0.15  CI95%: [-0.24, 0.50]  BF10: 0.31 | r: <-0.01  CI95%: [-0.25, 0.24]  BF10: 0.16 |
| BMI | r: -0.27  CI95%: [-0.59, 0.12]  BF10: 0.60 | r: -0.25  CI95%: [-0.47, 0.01]  BF10: 0.96 |
| *Abbreviations: AN: Anorexia Nervosa; BF10: Bayes Factor; BMI: Body Mass Index; CI95%: Confidence Interval; EDE-Q: Eating Disorders Examination- Questionnaire; HC: Healthy Controls.* | | |
